# Contrasting defensive strategies underlie differential susceptibility of corals to crown-of-thorns sea star (CoTS; *Acanthaster* cf. *solaris*) predation

**DOI:** 10.64898/2026.07.08.737165

**Authors:** Lucy M. Gorman, Stella L. Caon, Ariana S. Huffmyer, Maria Byrne, Sebastien Dutertre, Hollie M. Putnam, Suzanne C. Mills

**Affiliations:** PSL Université Paris: EPHE-UPVD-CNRS, UAR 3278 CRIOBE, BP 1013, 98729 Papetoai, Mo’orea, French Polynesia; Laboratoire d’Excellence “CORAIL”, France; Biological Sciences, University of Rhode Island, Kingston, RI 02881; School of Aquatic and Fisheries Sciences, University of Washington, Seattle, WA 98195; Marine Invertebrate Futures Group. School of Life and Environmental Sciences, Marine Science Institute, The University of Sydney, New South Wales, Australia; IBMM, Université de Montpellier CNRS, ENSCM, 34095 Montpellier, France; Institut Universitaire de France

**Keywords:** coral venom, toxins, crown-of-thorns sea star, pore-forming toxins, neurotoxins, metalloproteinases, *Acanthaster* cf. *solaris*, inducible and constitutive defences

## Abstract

Crown-of-thorns sea star (CoTS), *Acanthaster* cf. *solaris*, outbreaks are a major cause of hard coral cover decline across the west Pacific, threatening coral reefs. Coral taxa vary in susceptibility to CoTS predation from preferred (*Acropora* spp.) to non-preferred (*Porites* spp.), yet the mechanisms underlying these differences are poorly understood. We investigated coral defenses during an ongoing CoTS outbreak in Mo’orea, French Polynesia by examining gene expression (including putative toxin genes) in healthy and actively predated colonies of a preferred (*Acropora hyacinthus*) and a non-preferred (*Porites* sp.) coral prey species. During predation, *A. hyacinthus* exhibited molecular signatures of cellular stress responses involving oxidative stress signalling, inflammation, and tissue proteolysis. In contrast, *Porites* sp. showed enrichment of genes involved in mitochondrial metabolic adjustment and aerobic metabolism, suggesting metabolic compensation to maintain cellular function. Furthermore, *A. hyacinthus* demonstrated a reactive defense behaviour by differentially expressing toxins (e.g., kunitz-type neurotoxins) while *Porites* sp. employed constitutive expression of all putative toxins regardless of active predation, suggesting a proactive defense strategy. Together, these findings suggest that preferred and non-preferred coral prey exhibit fundamentally different molecular and defensive strategies during CoTS predation, shedding light on the evolutionary arms race between corals and their predators.

## 1. Introduction

Natural selection has generated a range of mechanisms across plants and animals allowing them to be effective at prey capture (predatory mechanisms) and resist predation themselves (defense mechanisms). One biological system that shows convergent evolution across the animal kingdom and used for ∼11 distinct functions is the evolution of venom systems (Casewell *et al*., 2013; Schendel *et al*., 2019). Venoms are mainly composed of toxins of protein/peptide origin, enzymes and neurotransmitters; allowing venomous organisms to disrupt the physiological homeostasis of their victim (Casewell *et al*., 2013).

Cnidarian venom systems are formed by specifically-evolved venom producing cnidocytes, cells that contain nematocyst vesicles that synthesise venom (Beckmann & Özbek, 2012). Upon mechanical or chemical stimulus of the hair-like cnidocil projection, the nematocysts fire a harpoon-like structure known as a thread into the tissue of the victim and release the venom (Beckmann & Özbek, 2012). Coral nematocysts are known to aid in prey capture and defense against corallivores (Lewis & Price, 1975; Gochfeld, 2004). However, unlike their jellyfish and sea anemone relatives, the composition and expression of coral venoms is relatively understudied. Coral venoms are known to contain pore-forming toxins, neurotoxins and toxic enzymes such as phospholipase A2s and metalloproteinases (Schmidt *et al*., 2019). However, it is unknown whether such defences can be specifically induced in response to predation by their major predator, the crown-of-thorns sea star (CoTS).

Population outbreaks of CoTS, can reach 1,500 sea stars per hectare (Dumas *et al*., 2016), and are a major cause of hard coral cover decline across the Indo-Pacific (De’Ath *et al*., 2012; Pérez-Rosales *et al*., 2021; Foo *et al*., 2024). One individual CoTS can consume around 5-12m^2^ of coral tissue per year (Chesher, 1969; Pearson & Endean, 1969) and can decimate 90% of corals on a single reef (Kayal *et al*., 2012; Leray *et al*., 2012). Worryingly, a recent study shows that despite coral cover reducing by 49% on Lizard Island, Great Barrier Reef, due to bleaching mortality and cyclone impacts, the density of CoTS increased by >96% (Levering *et al*., 2026). Furthermore, climate-driven poleward range expansion of CoTS to subtropical reefs in allowing CoTS larvae to be transported further afield (Nimbs *et al*., 2025; Sommer *et al*., 2025).

CoTS show a clear feeding preference for *Acropora* and pocilloporid corals (Keesing *et al*., 2019, 2021; Foo *et al*., 2024; Millican *et al*., 2024). During an outbreak CoTS displayed homing behaviour being attracted to *Acropora* when the abundance of this coral was > 33% (Ling et al., 2020). In contrast, these sea stars avoid massive coral genera such as *Porites* species (Millican *et al*., 2024), usually only feeding on this species when no other coral prey are available (Keesing *et al*., 2019). These feeding preferences between *Acropora* and *Porites* species are influenced by both nutritional content and amount of coral tissue available, as well as previous exposure to coral prey, with the latter being important for juvenile CoTS (Johansson *et al*., 2016; Keesing, 2021; Chandler *et al*., 2025).

CoTS feeding preference may also be influenced by several factors including coral venom and toxins (Barnes *et al*., 1970; Ormond *et al*., 1976; Deaker *et al*., 2021; Gorman et al., 2026). Corals defenses (e.g., nematocysts) can elicit lethal and sub-lethal injuries in naïve juvenile CoTS (not previously exposed to coral) (Yamaguchi, 1974; Deaker et al., 2021) and CoTS with sub-lethal injuries have been shown to revert back to herbivory to recover (Deaker *et al*., 2021). The lower susceptibility of CoTS predation for *Porites* relative to *Acropora* could reflect the differences in prey resource allocation, whereby slow-growing, stress-tolerant massive corals invest more heavily in tissue maintenance, immune function, and potentially defensive traits, whereas fast-growing *Acropora* species prioritize growth and competitive dominance (Darling *et al*., 2012). Understanding feeding preferences of CoTS is imperative to effectively predict the risk a CoTS outbreak poses to coral reef persistence and implement proactive management strategies.

Using bioinformatic analyses of *Acropora* and *Porites* species genomes we have shown that *Porites* spp., the non-preferred prey of CoTS, harbour different putative coral toxins and venom components to that of preferred coral prey, *Acropora* spp. (Gorman *et al*., 2026). Neurotoxins and pore-forming toxins were found in both taxa, but only *Porites* spp. had jellyfish toxins (CFXs), whereas only *Acropora* species harboured aerolysin-like toxins. Although toxin repertoires vary between species, it is not known whether these putative coral toxins are induced during predation by CoTS and how molecular responses to predation vary between preferred and non-preferred coral prey. Therefore, we quantified coral expression of defensive responses, including toxins, in corals in response to CoTS predation in non-preferred (*Porites* sp.) and preferred prey (*Acropora hyacinthus*) species, during a CoTS outbreak in Mo’orea, French Polynesia.

## 2. Methods

### 2.1. Specimen collection for transcriptomics

*Porites* sp. were collected from the lagoon at Ta’ahiamanu (−17.495210°, −149.851340°), Mo’orea, French Polynesia, across a 6-day period (30 January - 4 February 2024), between 9am-12pm. Similarly, *Acropora hyacinthus* specimens were collected from the outer reef (due to no individuals being eaten in the lagoon) adjacent to the Plage publique de Temae (−17.501364,-149.756490), Mo’orea between 12 March - 24 April 2024.

*A. hyacinthus* and *Porites* sp. colonies that had been actively preyed on by CoTS (“eaten”) and colonies that had not been preyed on (“control”) were chosen with *n* = 15 *per* species *per* predation status (*n* = 30 *per* species; **Fig. 1**). Control colonies were selected based on colonies that were not being actively eaten by CoTS and showed no observable historic CoTS feeding scars (**Fig. 1A, C**). Active predation was determined if a CoTS had its stomach everted over the colony, as confirmed by inverting the CoTS (**Fig. 2**). For the colonies being eaten, coral samples were excised from where the CoTS had been in contact with the colony. Coral biopsies were taken with stainless steel coral bone cutters, cleaned by wiping in seawater and rinsing with ethanol. The biopsies were placed in 1.5 mL screwcap tubes containing 0.75 mL of DNA/RNA shield (Zymo, Cat number: R1100-50). Colonies (eaten and control) were sampled at the same times (09:00-12:00) successively, alternating between an eaten and control sample. A subset of samples collected were utilized for molecular analyses described below (*n* = 8 *per* treatment *per* species).

**Figure 1.**
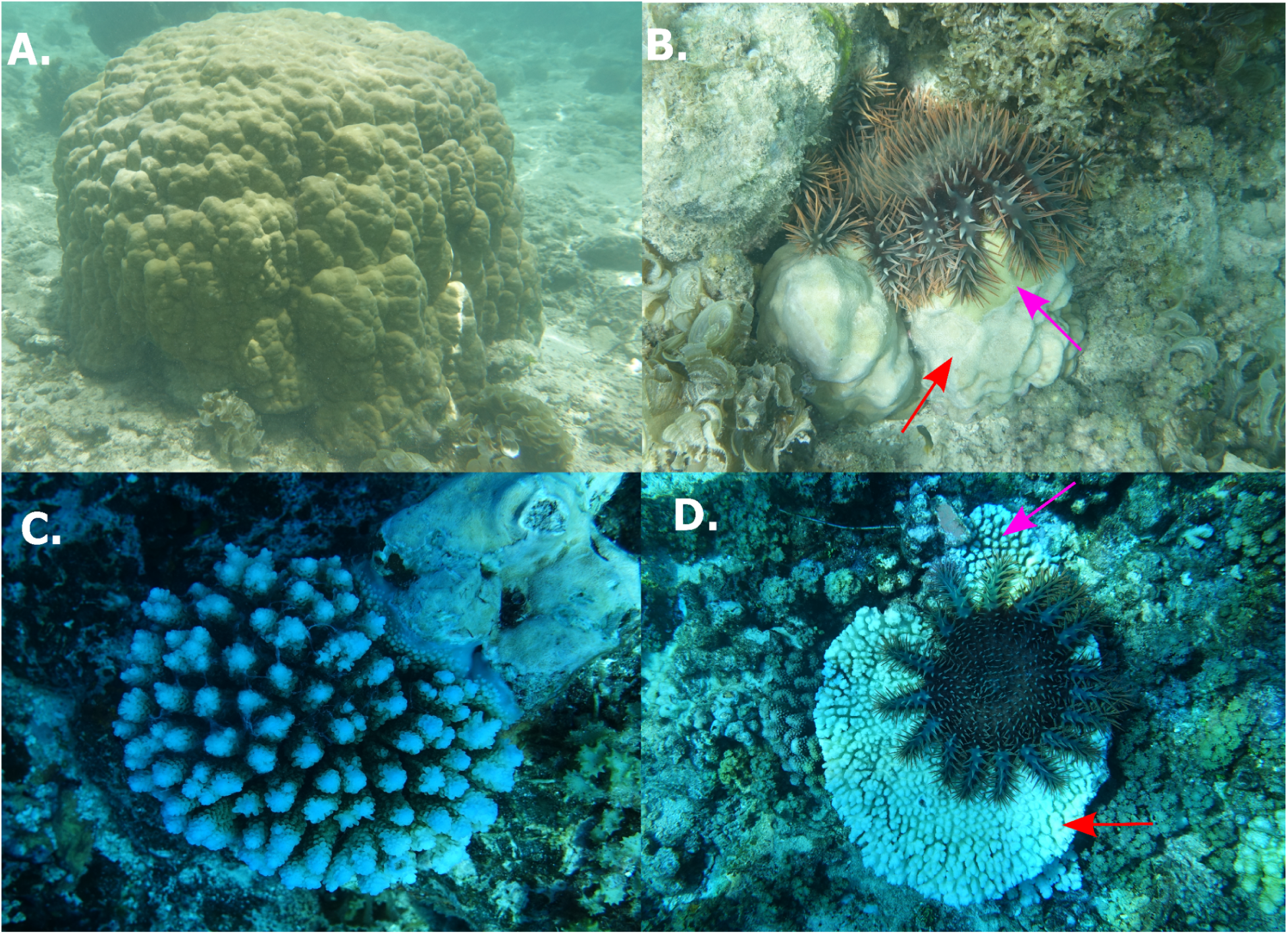
*Porites* sp. (**A**, **B**) and *Acropora hyacinthus* (**C**, **D**) during the 2022-2026 CoTS outbreak in Mo’orea, French Polynesia. Control colonies for each species were chosen based on no observable CoTS feeding scars (**A**, **C**). Colonies being eaten by CoTS (**B**, **D**) were chosen that were actively being consumed by CoTS during sampling and had an observable feeding scar (skeleton devoid of coral tissue (**red arrows**)) and areas adjacent to the scar with live coral tissue (**pink arrows**).

**Figure 2.**
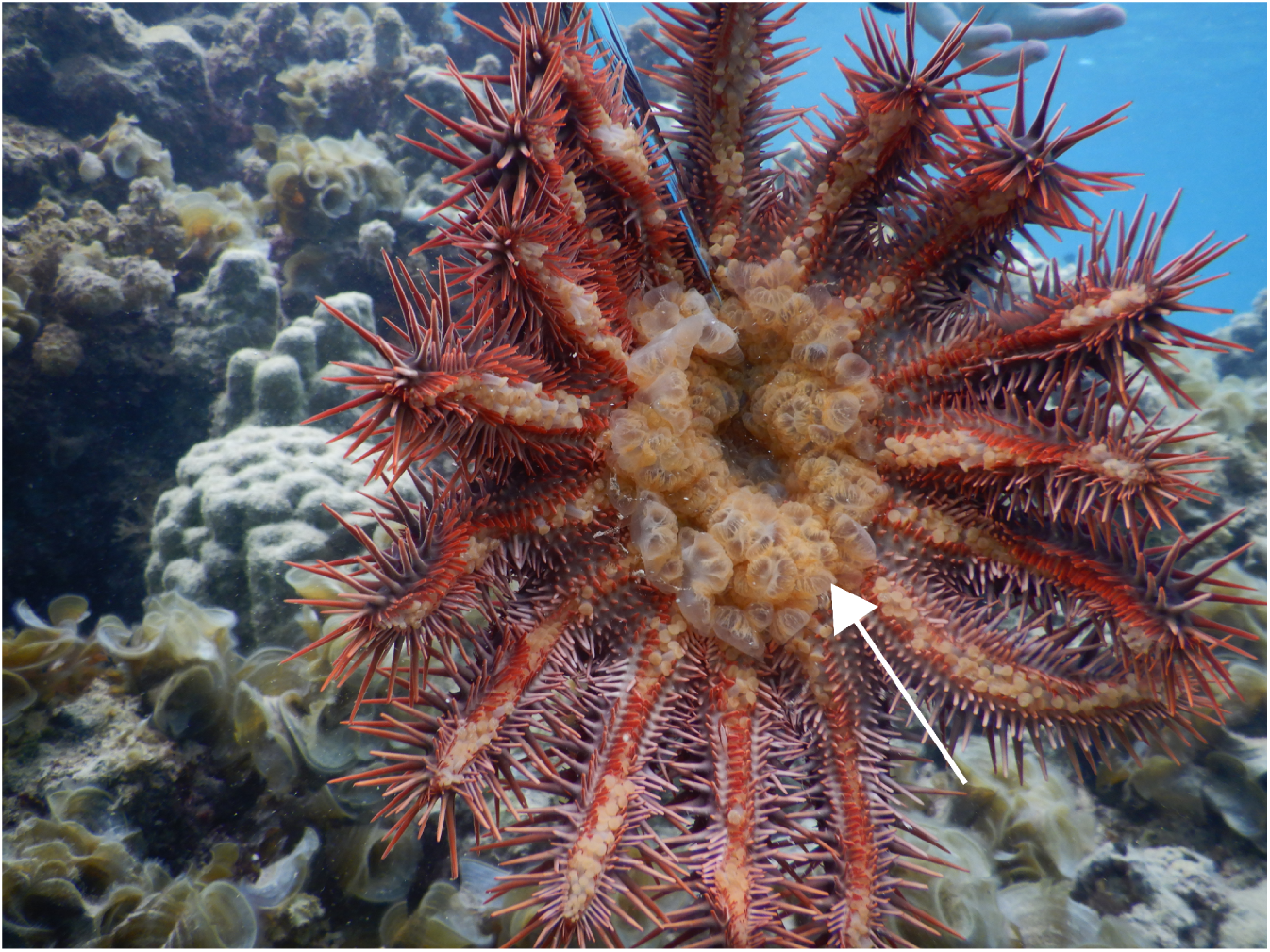
Crown-of-thorns sea star removed from a *Porites* sp. coral colony during feeding. White arrow shows its everted stomach.

### 2.2. RNA extraction

*Acropora hyacinthus* tissue (*n* = 8 *per* treatment) was removed from the skeleton by the preservation buffer and mixed by vortexing. *Porites* sp. tissue (*n* = 8 *per* treatment) remained on the coral skeleton while in the sampling buffer and the sample was therefore first pulverised with a metal blade and then mixed by vortexing. RNA was extracted from aliquots from each sample (300-750 µL) using the Zymo RNA/ DNA Quick Prep Mini kit (Cat number: D7001) following manufacturers instructions and quality examined using an N60 nanospectrophotometer (IMPLEN, Munich, Germany) and Bioanalyser (Agilent, Santa Clara, California).

### 2.3. RNA sequencing and library construction

RNA sequencing and library construction were carried out at MGX-Montpellier GenomiX, University of Montpellier, CNRS, INSERM, with ribosomal RNA depletion using Illumina Ribo-Zero (Cat number: 20040526). Libraries were prepared *via* polyadenylated RNA selection with polyT capture beads using Illumina’s Truseq stranded mRNA kit (Cat number: 20020594) according to manufacturer’s instructions. Libraries were validated by DNA quantification for fragment size and concentration using a Fragment Bioanalyzer (Agilent) and qPCR (Roche), respectively. The resulting RNA was sequenced 2 x 150 bp on one lane of a Novaseq S4 (Illumina, San Diego, California).

### 2.4. Bioinformatics

One sample of pair-end reads in the *Porites* sp. eaten treatment had an abnormally high number of reads in the sequence library (> 610 million) and seqTK software v.1.4 (Li, 2023) was used to randomly subset to 90 million reads (average number of reads in all other samples). Paired-end reads from all samples were then filtered and trimmed using fastp v.0.23.2. (Chen, 2023) with the following parameters: a qualified base quality value of 20; an unqualified base limit of 10%; window size of 5 and a quality threshold of 20 for the window. Reads were assessed for quality using fastQC (Babraham Institute, 2023) and multiQC (v.1.12, Ewels *et al*., 2016). *Acropora hyacinthus* samples were mapped to the *A. hyacinthus* genome (Lopez-Nadam *et al*., 2023) using STAR (v.2.7.11.b, Dobin *et al*., 2013). *Porites* cannot be reliably identified to species visually (Forsman *et al*., 2009), therefore *Porites* spp. samples were mapped to both an Australian *P. lutea* genome (Robbins *et al*., 2019) and a French Polynesia *P. evermanni* genome (Planes *et al*., 2019). All samples from *Porites* species showed higher mapping to the *P. evermanni* genome and therefore, the *P. evermanni* genome was chosen for alignment using STAR (v.2.7.11.b).

A gene count matrix was generated using StringTie v3.0.0 (Shumate *et al.,* 2022). Genes with 0 counts were then removed from the dataset and filtered using pOverA (genefilter R package; Gentleman *et al*., 2026). Outliers were identified using Principle Coordinate Analysis (PCA) plots of expression of all genes detected. Samples with a low percent of total mapped reads (<5%) were removed from the datasets as well as those that were outliers on PCA plots (determined *via* visual inspection), totalling one eaten *A. hyacinthus* and four eaten *Porites* sp. replicates being removed from further analysis (**Table S1**). This resulted in a sample size of *n* = 7 and *n* = 8 eaten and control *A. hyacinthus*, respectively and *n* = 4 and *n* = 8 eaten and control *Porites* sp., respectively. Overall, genes were retained that were present in 44% of samples in *A. hyacinthus* (proportion of samples in smallest treatment group) and 34% of samples in *Porites sp.* with transcript counts >10, resulting in 18,258 total genes for *A. hyacinthus and* 22,214 for *Porites* spp.

### 2.5. Analysis

Genes differentially expressed in “eaten” colonies were identified using differential gene expression analysis. Gene counts were normalized using a variance stabilized transformation in the DESeq2 package in R (Love *et al*., 2014). A permutational analysis of variance (PERMANOVA) was then used to test for the effects of predation status for all genes that passed filtering thresholds followed by a permutational analysis of dispersion (PERMDISP) to test for variation in dispersion in the vegan package in R (Oksanen *et al*., 2025). A Wald test in DESeq2 was then used to identify differentially expressed genes between eaten and control colonies within each species. Differentially expressed genes (DEGs) were identified as genes with false discovery rate (FDR) adjusted *P* < 0.05.

The general feature format (gff) file of the *A. hyacinthus* genome was formatted and cleaned using the AGAT toolkit v1.4.1 plugin in python (Dainat, 2022). Functional enrichment of the *A. hyacinthus* genome was performed as in Conn *et al*. (2025) using funannotate v.1.8.17 (Palmer & Stajich, 2020). Briefly, the *A. hyacinthus* genome was soft-masked using the mask function in funannotate v.1.8.17 (Palmer & Stajich, 2020). The soft-masked *A. hyacinthus* genome was then annotated by firstly assigning gene function and ontology using the eggnog mapper v.2.1.12 (Cantalapiedra *et al*., 2021) and InterProScan v.5.73-104.0 (Jones *et al*., 2014). The outputs of these were then inputted into funannotate wrapper to run alongside funannotate. Funannotate then searches the PFAM (Paysan-Lafosse *et al.*, 2025), CAZy (Drula et al., 2022), UniProt (The UniProt Consortium, 2025) and GO (Ashburner *et al*., 2000; The Gene Ontology Consortium, 2026) databases and assigns all the results to each gene. The Blast2GO suite (Götz *et al*., 2008) in OmicsBox (v3.5; OmicsBox, 2019) was used to functionally annotate the *P. evermanni* genome. Functional enrichment was then conducted using the topGO package in R (Alexa & Rahnenfuhrer, 2025). A node size of 5 was chosen with an ontology of biological process (“BP”) and an annotation setting of annFUN.gene2GO. A weighted Fisher exact test with *P* < 0.05 was used to identify significant GO terms in DEGs for each species. Functional annotations for *A. hyacinthus* and *Porites* sp. genomes can be found in the GitHub repository (see *Data Accessibility*).

### 2.6. Toxin expression: NCBI BLAST+ and structural comparisons

In order to quantify the number of putative toxin genes expressed, both the differentially expressed genes (DEGs) and all the expressed genes (EGs) of each species (18,258 genes in *A. hyacinthus* and 22,214 in *Porites* sp.) were searched against the cnidarian filtered UniProt ToxProt database (342 proteins; accessed December 2025) and the database we had previously created (“custom database”; Gorman *et al.,* 2026) using the NCBI BLAST+ command line. We only collected the top hit (highest query cover) of the BLAST against the custom database due to the high number of paralogs which would be returned. Parameters for the BLAST search against the UniProt ToxProt were: an e-value of 1e-5, a word size of 6, a gapopen size of 11, a gap extend value of 1, a query coverage of 70% and the option to filter the query sequence with SEG was disabled. These parameters were repeated for the custom database searches except the e-value was 1e-10 and the query coverage was 80% to reduce false-positive hits. Hits to EGs and DEGs were then cross checked between the two databases, resulting in 50 hits for *A. hyacinthus* and 55 for *Porites* sp. to putative toxins across their expressed gene pools (**Table S2, 4**). Similar 3D protein structures were determined by searching the amino acid sequence of putative toxins against the Alphafold server (Fleming *et al*., 2025) and the toxin returned with the highest 3D structural similarity (pLDDT score) from Alphafold was recorded (**Tables S2-5**). One gene (*Peve_00044641*) had an unknown amino acid residue “X” at position 29 in the sequence that was substituted for an alanine as a placeholder to predict 3D structure with AlphaFold (**Table S3)**. Another gene (*Peve_00015061*) had several unknown amino acid residues “X”, therefore its 3D structure was not predicted with AlphaFold (**Table S3**). Hits which were returned with structural 3D similarity to non-venomous proteins were removed from analysis. This left a total of 47 hits for *A. hyacinthus* and 52 for *Porites* sp. to putative toxins (**Table 1**).

**Table 1.** Putative toxin gene expression during a CoTS outbreak in *Acropora hyacinthus* and *Porites* sp. colonies. Numbers in parentheses refer to metalloproteases or kunitz-type neurotoxin proteins with strong candidacy of functioning in envenomation.

|  | Toxin | *Acropora hyacinthus* | *Porites* sp. |
| --- | --- | --- | --- |
| Differentially expressed genes (DEGs) between eaten and control colonies | Metalloproteinase | 4 (0) | 0 |
|  | Kunitz-type toxin | 1 (1) | 0 |
|  | Insulin-like peptide | 1 | 0 |
| Total toxin genes expressed (genes expressed in both eaten and control colonies; TGs) | Metalloproteinase | 21 (2) | 30 (7) |
|  | Kunitz-type toxin | 9 (3) | 12 (7) |
|  | Insulin-like peptide | 2 | 4 |
|  | NEP3-like neurotoxin | 1 | 0 |
|  | PLA2 | 3 | 1 |
|  | CFX-like | 0 | 2 |
|  | Actinoporin | 1 | 1 |
|  | MAC-PF | 2 | 1 |
| | SCRiPS-δ | 2 | 1 |
| Total |  | 47 | 52 |

### 2.7. Identification of putative metalloproteinases and kunitz-type neurotoxins

As metalloproteinases and proteins with the Kunitz-type BPTI domain (found in kunitz-type neurotoxins) can also function in non-venomous tissue, the putative toxins with the highest candidacy of functioning in envenomation are highlighted in **Table 1** (see **Supplemental file 1** for methodological criteria for defining strong candidacy). Briefly, a putative metalloproteinase was referred to as a strong candidate for venom functioning if it showed sequence and structural homology to nematocyst expressed protein (NEP) 6 from *Nematostella vectensis*, hypothesised to be a toxic metalloproteinase (Moran *et al*., 2013; **Tables S6**). Similarly, a kuntiz-type neurotoxin was referred to as a strong candidate if they showed: a structural similarity to actitoxins/stichotoxins in anthozoans (pLDDT > 86) or a structural homology to the thrombin inhibitor, hemalin, from *Haemaphysalis longicornis* (pLDDT = 91.38); and additionally, a sequence homology to the kunitz-type neurotoxins PcKuz1 and PcKuz3 in the zoanthid *Palythoa caribaeorum* (Liao et al., 2018; **Tables S7**).

### 2.8. 3D Protein Structure Prediction of Chironex fleckeri-like toxins (CFXs)

Due to their lower query cover to true CFXs, putative CFX-like toxins from *Porites* sp. had their 3D structure predicted using ColabFold v1.6.1 (Mirdita et al., 2022) with default parameters. Comparison of 3D protein structures between putative CFX-like toxins from *Porites* sp. and the toxins returned with the highest 3D structural similarity from FoldSeek (Van Kempen et al., 2024) in the AFDB-SwissProt was then conducted in DALI (Holm et al., 2022). 3D structures were also compared and visualised in ChimeraX v1.12 (Meng et al., 2023) using the matchmaker program with default parameters (**Table S8**). Regions with high candidacy of an amphipathic helix region (which contributes to pore formation in PFTs) were observed with sequence searches using HELIQUEST v2.0 (Gautier et al., 2008).

## 3. Results

### 3.1. Divergent molecular responses to predation in preferred and non-preferred coral species

CoTS feeding influenced gene expression in *A. hyacinthus* (PERMANOVA, *P* = 0.002; **Fig. 3A**), with 1,405 genes differentially expressed (DEGs) in response to predation (**Fig. 3B, C**). DEGs in *A. hyacinthus* were enriched for 38 significant (**Fig. S1**) gene ontology (GO) terms (652 significant genes out of 8,095 genes with annotation information) and were strongly enriched for superoxide anion generation (GO:0042554), response to oxidative stress (GO:0006979), NADPH regeneration (GO:0006740) and mitochondrial electron transport (GO:0006120) (**Fig. 3D & S1**). Terms related to metabolic stress and energy metabolism were also enriched in DEGs including proteolysis (GO:0006508), collagen catabolic process (GO:0030574), unsaturated fatty acid biosynthetic process (GO:0006636) and glucose homeostasis (GO:0042593) (**Fig. 3D & S1**). Cell signaling and inflammation were also affected by predation indicated by enrichment in non-canonical and canonical Wnt signaling pathways (GO:0035567 and GO:0060070), Rho protein signal transduction (GO:0007266), and regulation of kinase activity (GO:0033674), including NF-kappaB (GO:0051092) (**Fig. 3D & S1**).

**Figure 3.**
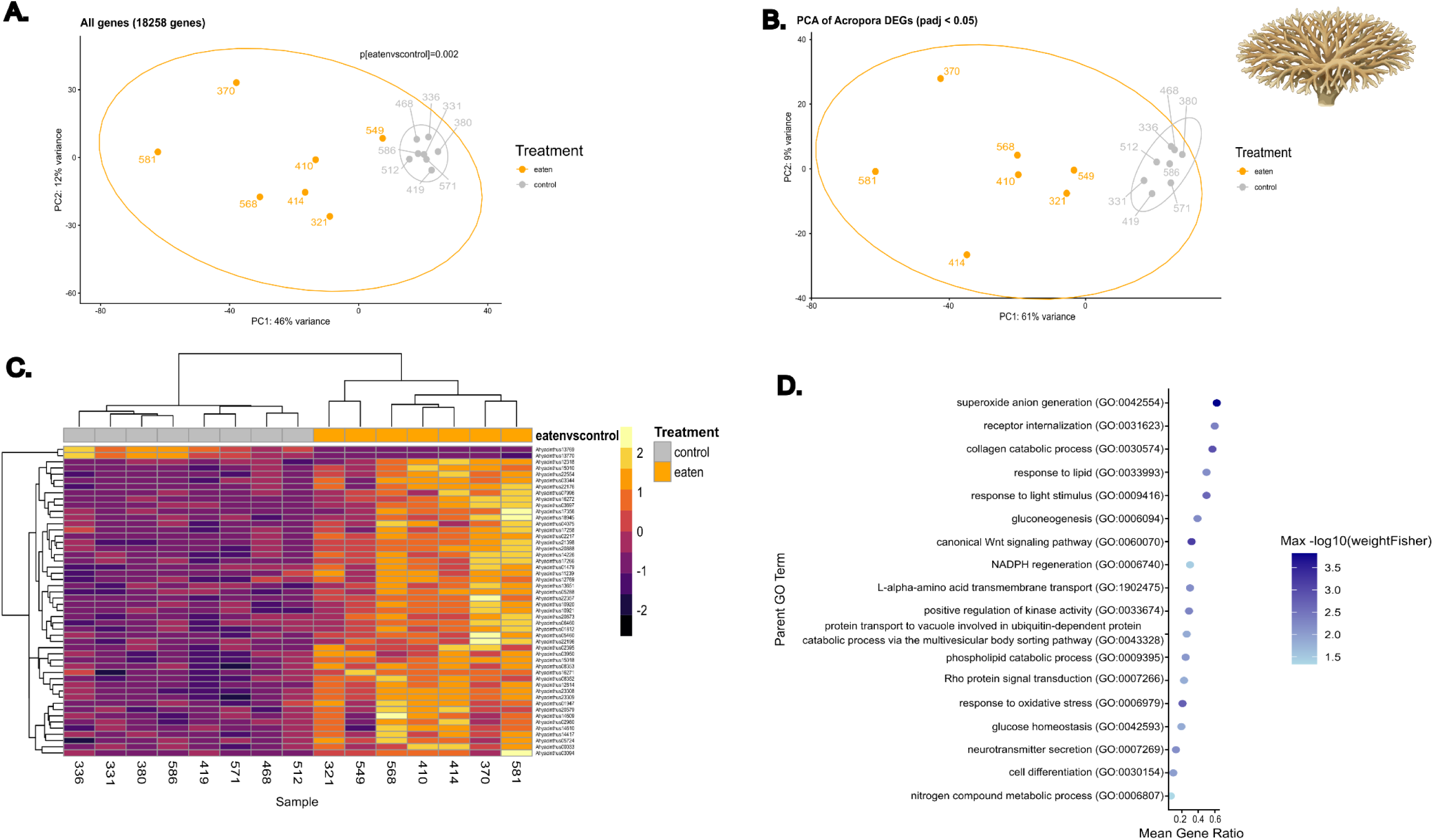
Gene expression data from eaten (*n* = 7) and control (*n* = 8) *Acropora hyacinthus* colonies during a CoTS outbreak. **A.** Principle Coordinate Analysis (PCA) of VST-transformed global gene expression. **B.** PCA of DEGs (*n* = 1405). **C.** Heatmap of top 50 DEGs. **D.** Significantly enriched Gene Ontology (GO) biological process (BP) parent terms (*n* = 18; adjusted *p* < 0.05) and Mean Gene Ratio (x axis, dot size) representing the proportion of differentially expressed genes (DEGs) in each category relative to the total genes in that category. Color indicates weighted Fisher test value.

In contrast to *A. hyacinthus*, in *Porites* sp. CoTS feeding did not significantly influence gene global expression (perMANOVA, *P* = 0.57; **Fig. 4A**). However, there were 18 differentially expressed genes between eaten and control colonies (**Fig. 4B, D**). These genes were enriched for 74 significant (**Fig. S2**) gene ontology (GO) terms (6 significant genes out of 13,991 genes with annotation information) (**Fig. 4C**). *Porites* sp. colonies being actively eaten by CoTS showed enrichment in genes related to DNA repair, mitochondrial metabolic adjustment and efficient aerobic metabolism. Specifically, DEGs were enriched for GO terms including mitochondrial electron transport, cytochrome c to oxygen (GO:0006123), respiratory chain complex IV assembly (GO:0008535), oxidative phosphorylation (GO:0006119), and ATP metabolic process (GO:0046034) (**Fig. 4C & S2**). The strongest gene ratio GO terms were for ectodermal digestive tract morphogenesis (GO:0048567) and depyrimidination (GO:0045008). In addition, eaten *Porites* sp. colonies exhibited differential expression in genes related to stored energy metabolism including heme biosynthetic process (GO:0006783) and fatty acid beta-oxidation using acyl-CoA (GO:0033539).

**Figure 4.**
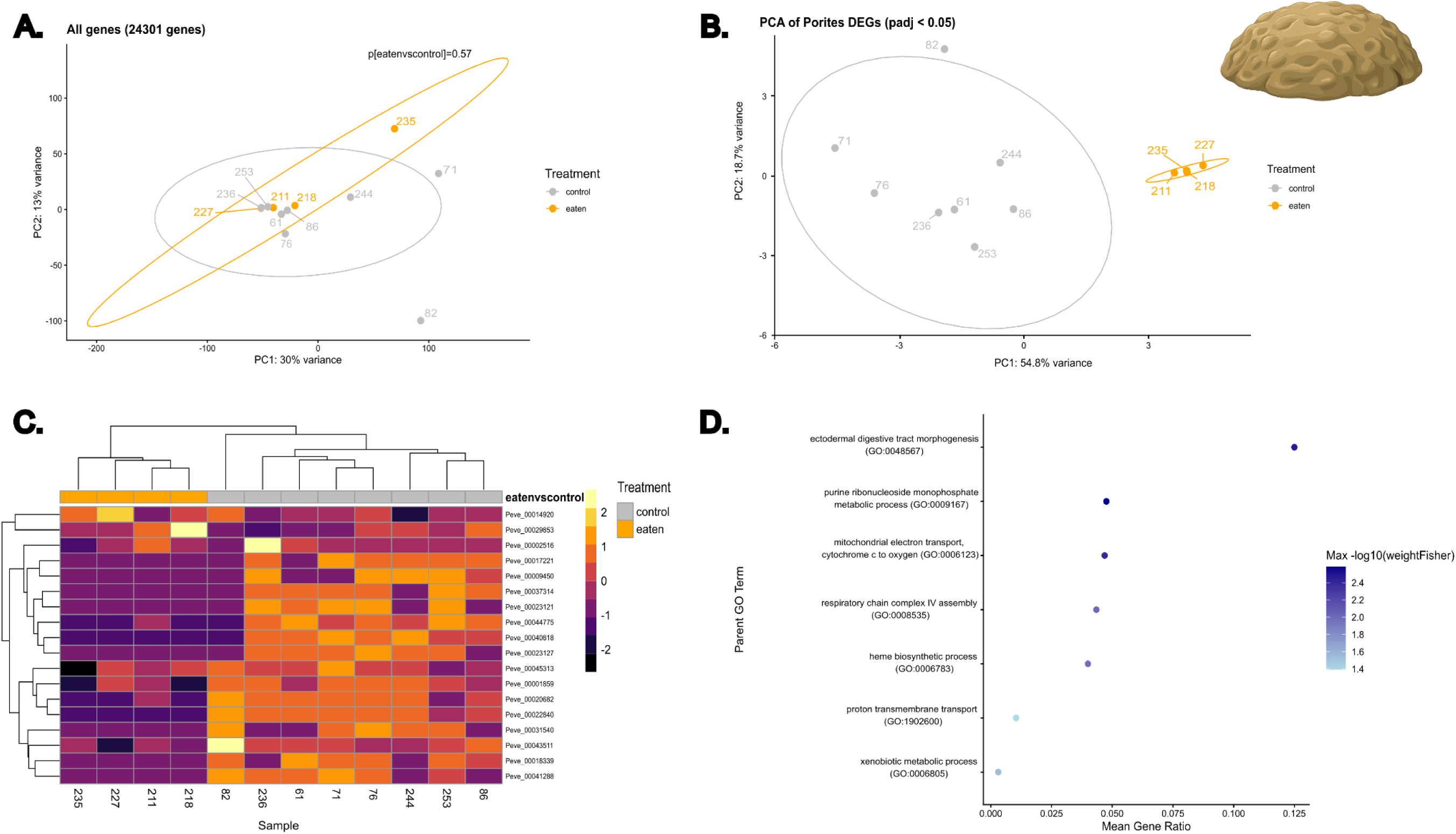
Gene expression data from eaten (*n* = 4) and control (*n* = 8) *Porites* sp. colonies during a CoTS outbreak. **A.** Principle Coordinate Analysis (PCA) of VST-transformed global gene expression. **B.** PCA of DEGs (*n* = 18). **C.** Heatmap of DEGs. **D.** Significantly enriched Gene Ontology (GO) biological process (BP) parent terms (*n* = 14; adjusted *p* < 0.05) and Mean Gene Ratio (x axis, dot size) representing the proportion of differentially expressed genes (DEGs) in each category relative to the total genes in that category. Color indicates weighted Fisher test value.

### 3.2. Toxin expression

We investigated expression of toxin genes (TGs) (genes identified as toxins expressed across all colonies), as well as differentially expressed toxins (DEGs identified as toxins) in *A. hyacinthus* and *Porites* sp. A total of 47 and 52 putative TGs corresponding to neurotoxins, pore-forming toxins and toxic enzymes were expressed overall in both control and eaten colonies in *A. hyacinthus* and *Porites* sp., respectively (**Table 1, S2 & S3**). *A. hyacinthus* colonies showed 1,405 DEGs, with only six corresponding to putative toxin genes - four metalloproteinases, one kunitz-type neurotoxin and one insulin-like peptide (**Table 1**). Of the six putative toxin DEGs in *A. hyacinthus*, all except the insulin-like peptide showed higher expression in the eaten colonies compared to the control (**Table S4**). Interestingly, *Porites* sp. colonies showed no differential expression of toxin genes in response to CoTS predation (**Table 1**). Of the TGs, *A. hyacinthus* and *Porites* sp. expressed genes from similar toxin families with metalloproteinases being the most abundant toxin family, followed by Kunitz-type toxins and SCRiPS neurotoxin homologs (**Tables 1, S4, S5**).

*A. hyacinthus* and *Porites* sp. both expressed genes corresponding to the pore-forming toxins (PFTs) DELTA-actitoxin family actinoporins and membrane attack complex/perforin (MAC-PFs) PFTs in anthozoans; insulin-like peptides; the neurotoxic small cysteine-rich protein (SCRiP)*-δ;* and, toxic phospholipase A2s (**Table S2-S5**). PFTs (actinoporins, MACPFs, and CFX-like toxins in *Porites* sp.) were not as highly expressed as metalloproteinases or neurotoxins (**Table S4 & S5**). Structural similarities revealed high similarities across PFTs and PLA2s in *A. hyacinthus* and *Porites* sp., with: actinoporin homologs from both species showing structural similarity to Fragaceatoxin E from *Actinia fragacea* (pLDDT = 97.56); MAC-PFs from both species showing structural similarity to the haemolytic PsTX-60B from *Phyllodiscus semoni* (pLDDT = 90.06; Satoh *et al*., 2007); and, phospholipase A2 genes from both species showing structural similarity to Phospholipase A2-actitoxin-Cgg2a from *Condylactis gigantea* (pLDDT = 96.19; **Table S2 & S3**).

*A. hyacinthus* expressed a unique putative neurotoxin with three Shk domain repeats whilst *Porites* sp. expressed CFX-like homologs (**Table 1**). The putative neurotoxin in *A. hyacinthus* (*Ahyacinthus20286*; **Table S2**) showed sequence homology to NEP3 from *Nematostella vectensis* and structural homology to both NEP4 and NEP8 from *Nematostella vectensis*. All these NEP proteins belong to the NEP3 toxin family (Columbus-Shenkar et al., 2018) that all contain three Shk domain repeats, similar to *Ahyacinthus20286.* Notably, only *Porites* species expressed genes (*Peve_00009963* and *Peve_00015288*) that matched to two *Porites* CFX homologs found in our previous bioinformatics study (Gorman et al., 2026), with *A. hyacinthus* not expressing any homologs to CFXs (**Table S3**). These findings agree with our bioinformatics study that showed the absence of CFX toxins across five *Acropora* species (Gorman et al., 2026). Although both *Porites* sp. genes met the criteria to the *Porites* CFX-like toxins, they did not meet the query coverage threshold to the true jellyfish CFXs (≥ 70%; **Table S3**). One of the putative CFX-like toxins (*Peve_00009963*) harboured the pesticidal crystal protein domain (IPR036716) found in Cry-like toxins (**Table S3**), and this domain has been reported to be the structurally closest toxin of true CFXs (Brinkman *et al*., 2014). Both CFX-like toxins (*Peve_00009963* and *Peve_00015288*) showed structural similarity to type II JFT-1b toxins (TM score **≥** 0.64; **Table S8**) and harboured amino acid residues with a strong candidacy of forming an amphipathic helix which contributes to pore formation in PFTs (Hong *et al*., 2002; **Supplemental file 1**). Both CFX-like toxins showed regions of structural conservation to *Chironex fleckeri* CfTX-1 in the transmembrane spanning region (**Supplemental file 1**). The two putative CFX-like toxins found in this current study were distinct from one another, only showing a 39% sequence identity, implying that they may belong to different clades of CFX-like PFTs.

## 4. Discussion

Coral reefs are under threat from multiple stressors with climate change driven mortality (bleaching, cyclones) being exacerbated by the damage caused by CoTS outbreaks (Garing *et al*., 2026). Understanding coral species resilience and resistance to CoTS predation is imperative to inform management strategies. We found differences in gene expression between preferred (*A. hyacinthus*) and non-preferred (*Porites* sp.) during CoTS predation, relating to variation in energetic and defensive strategies. A preferred species, *A. hyacinthus,* exhibited widespread systemic homeostasis dysregulation and was in a stress responsive state during CoTS predation, fueled by pathways involved in ROS oxidative stress signalling, inflammation and tissue proteolysis. In contrast, a non-preferred species, *Porites* sp., exhibited expression of genes involved in mitochondrial metabolism, ATP generation and aerobic metabolism to potentially displayed a reactive defense strategy to CoTS predation with inducible defences, indicated by differentially expressed putative toxins (e.g., metalloproteinases, an insulin-like peptide and a kunitz-type neurotoxin). *Porites* sp. exhibited a proactive defense strategy with constitutive expression of putative toxin genes. Together, these findings suggest that preferred and non-preferred coral prey exhibit different defensive strategies during crown-of-thorns sea star predation, providing insights into species-specific predation responses.

The difference in defence strategies between *A. hyacinthus* and *Porites* sp. may reflect underlying differences in their metabolic and life-history traits (i.e., slow-growing *Porites* vs fast-growing *Acropora*) and the amount of energy they can allocate to toxin production. Toxin expression is energetically costly, and venomous organisms are shown to employ venom frugality *via* “venom metering”, where venom production is tailored for specific instances and organisms, reducing unnecessary energy expenditure (Morgenstern & King, 2013). In the anthozoan *Nematostella vectensis,* venom biosynthesis increases respiration rate and toxin genes are downregulated under abiotic stress to metabolically compensate (Sachkova *et al*., 2020). Massive species such as *Porites*, have higher metabolic and lower growth rates than branching species (Gates & Edmunds, 1999; Putnam *et al*., 2012). Thus, *Porites* sp. may be able to invest more metabolic resources overall in toxin production, allowing them to keep a continuous toxin expression regardless of CoTS predation state, making them more effective at defending themselves. In contrast, *Acropora* exhibit lower metabolic rates but higher growth rates and prioritise their energy budget towards growth and reproductive pathways (Gates & Edmunds, 1999; Darling *et al*., 2012; Putnam *et al*., 2012). These traits allow them to quickly occupy space and dominate reefs following disturbance, but make them more vulnerable to stressors (Loya *et al*., 2001; Keesing *et al*., 2019). This trade-off between toxin production and growth rates has been observed in *Nematostella vectensis*, where reduced expression of the neurotoxin Nv1 facilitated faster growth rates and increased reproduction (Surm *et al*., 2024).

It should also be noted that *A. hyacinthus* samples were collected closer to the time of maximum annual sea temperature *versus Porites* sp. These differences in collection time may have contributed to *A. hyacinthus* allocating more resources to stress responses (possibly including thermal sensitivity) and less resources to toxin production. Indeed, it is well noted that thermal stress increases the respiration and metabolic rate of corals (Coles & Jokiel, 1977; Parry *et al*., 2025). In the context of CoTS predation, this energy budgeting may make *A. hyacinthus* employ a more reactive toxin expression during predation compared to *Porites* species, potentially influencing their susceptibility to CoTS predation.

Although overall *A. hyacinthus* and *Porites* sp. displayed a more reactive and proactive behaviour respectively, both species did express a subset of toxins constitutively regardless of their predation status. This could reflect that neighbouring corals not being eaten CoTS may be able to upregulate their toxin defence arsenal prior to CoTS predation in response to semiochemical cues released by CoTS and/or the coral colony being eaten. Indeed, both CoTS and corals are known to release and respond to semiochemical cues (Armoza-Zvuloni *et al*., 2016; Webb *et al*., 2024), and the chemical cues released by CoTS feeding on coral have been shown to deter coral reef fish (Coppock et al., 2016), emphasising that this predatory behaviour likely releases infochemicals that coral reef organisms (including neighbouring corals) can sense. Another factor that may contribute to the constitutive expression of certain toxins is that corals may employ a constant basal level of toxin expression to allow them to be effective at heterotrophy and prey capture. It should be noted however that cnidarians have been shown to harbour different toxins for defense *versus* prey capture (Columbus-Shenkar *et al*., 2018). Whether corals can increase their toxin expression to constitutively express toxins to allow for effective heterotrophy, or can rapidly deploy their toxin repertoire by sensing chemicals released by CoTS and/or eaten neighbours, warrants future investigation.

The majority of toxin families expressed across *A. hyacinthus* and *Porites* sp. were the same with the dominant families being metalloproteinases and neurotoxins, as seen in our bioinformatics study (Gorman et al., 2026) and studies on jellyfish venom (Li et al., 2022; Yang et al., 2024). However, two toxin families differed between *A. hyacinthus* and *Porites* sp. - a neurotoxic NEP3-like homolog only expressed in *A. hyacinthus* and two jellyfish CFX-like homologs only expressed in *Porites* sp. NEP3 family toxins in *N. vectensis* are expressed in different sub-populations of nematocysts and bioactivity assays conducted on zebrafish larvae with NEP3 showed neurotoxicity-type bioactivity (Columbus-Shenkar et al., 2018). NEP3 toxins have been shown to be lost in some hexacorallian lineages, but a homolog to the NEP3 toxin was found in the *Acropora digitifera* transcriptome (Columbus-Shenkar et al., 2018), with our current study further confirming the presence across other acroporid species. Both the putative CFX-like toxins in *Porites* sp. showed regions of structural conservation to *Chironex fleckeri* CfTX-1 in the transmembrane spanning region which is hypothesised to play a major role in the mode of action of CFXs (Andreosso et al., 2018). CFXs represent some of the most rapid and potent toxins in cnidarians, with type I CFXs causing cardiovascular collapse whilst type II cause haemolysis (Brinkman et al., 2014; Vega-Tamayo et al., 2025). The CFX-like toxins found in this current study differed from our previous bioinformatics study where CFX homologs found in other *Porites* species (*P. rus*, *P. compressa* and *P. australiensis*) showed high query coverage to true CFXs (≥ 70%; Gorman et al., 2026). However, the CFX-like toxins in the current study showed more divergence (< 70% query cover) to the true jellyfish CFXs.

The abundance of expression of putative kunitz-type neurotoxin genes in both *A. hyacinthus* and *Porites* sp. containing structural homology to actitoxins/stichotoxins is noteworthy, especially in light of recent evidence from scleractinians that these toxins may play a vital role in toxicity. For example, a recent study has shown that a kunitz-type neurotoxin from *Porites lutea,* PlKuz1, is a selectively potent inhibitor of potassium channels and is lethal to zebrafish (Chen et al., 2025a). Similarly, a kunitz-like toxin from the scleractinian *Goniopora columna* exerts anticoagulant effects (Chen et al., 2025b). One of the putative kunitz-type neurotoxin genes in *A. hyacinthus* was identified as a differentially expressed toxin during CoTS predation. These findings, in conjunction with a kunitz-type neurotoxin isolated from the CoTS predator *Charonia tritonis* degrading CoTS tube feet and paralysing mice (Zhang *et al*., 2022), makes kunitz-type neurotoxins in scleractinian venoms an intriguing avenue for future exploration.

In addition to metalloproteinases and kunitz-type neurotoxins, a diversity of other toxin families were found across both species including: actitoxin actinoporins, MAC-PFs; insulin-like peptides; the neurotoxic SCRiPS*-δ;* and, toxic phospholipase A2s. A putative SCRiPS neurotoxin was the most highly expressed toxin in eaten *Porites* sp. colonies, with SCRiPS*-δ* previously being hypothesised to be the dominant toxic SCRiPS family involved in scleractinian defence (Barroso et al., 2024). Both *A. hyacinthus and Porites sp.* had sequence homologs to haemolytic DELTA-actitoxins actinoporins - Bandaporin and Or-A, respectively (Il’ina et al., 2005; Kohno et al., 2009; Monastyrnaya *et al*., 2010). Genes homologous to DELTA-actitoxins have also been found in fire coral *Millepora complanata* (Hernández-Elizárraga et al., 2022) and scleractinians *Montipora capitata* (Helmkampf *et al*., 2018) and *Orbicella* species (Wright et al., 2019), implying that this class of actinoporins is widespread throughout cnidarians. In *M. complanata*, the increased abundance of the DELTA-actitoxins was correlated with an increase in the haemolytic activity of the proteome (Hernández-Elizárraga et al., 2019). Indeed, the function of SCRiPS neurotoxins and DELTA-actitoxins actinoporins in scleractinian venoms warrants further investigation.

Control and eaten colonies of *A. hyacinthus* and *Porites* species also expressed putative toxic PLA2 genes that had structural similarity to PLA2-actitoxin-Cgg2a from *Condylactis gigantea* that induces myotoxicity and edema in mice (Romero et al., 2010). *Acropora* elicits higher PLA2 activities than those recorded across Cubozoa and Scyphozoa jellyfish species, including box jellies (Nevalainen *et al*., 2004). Thus, the increased expression of toxic PLA2 homologs found in this current study may contribute to the higher levels of PLA2 activity observed in *Acropora* species. Likewise, insulin-like venom peptides were expressed in both *A. hyacinthus and Porites s*p., with one of these homologs differentially expressed in eaten *A. hyacinthus* colonies. Insulin from cone snail venom incapacitates prey by collapsing their blood sugar levels, rendering them paralysed (Safavi-Hemami *et al*., 2015). Insulin and insulin-like peptides are widespread in venomous animals including cnidarians (Laugesen *et al*., 2022; Delgado *et al*., 2024), however whether these peptides function as in cone snail venom is yet to be determined (Laugesen *et al*., 2022).

### Conclusion

Our data show that the preferred prey of CoTS (*A. hyacinthus*) in Moorea undergo widespread regulatory dysfunction, strong ROS stress and signalling responses, homeostatic instability and tissue damage during predation. Contrastingly, non-preferred prey (*Porites* sp.) compensate metabolically and enrich pathways involved in oxidative metabolism (mitochondrial electron transport and respiration), retaining overall cellular function. Moreover, *A. hyacinthus* colonies showed a reactive defensive behaviour to CoTS predation by upregulating several putative toxin genes. *Porites* sp. had a proactive strategy, evidenced by constitutive expression of toxin genes with a richer venom repertoire. In an ecological and coral reef restoration context, these findings suggest that *A. hyacinthus* is more susceptible to CoTS predation and less likely to invest in a full toxin repertoire for defence. Future studies should prioritise proteomics and bioactivity of these venom components, both individually and combined, to generate an accurate venom proteome for these species and enhance our knowledge of how they function against CoTS during predation (Walker *et al*., 2020). Furthermore, future advances in single-celled transcriptomics of scleractinians (Levy *et al*., 2021; Han *et al*., 2025) will help to identify which of these genes are found in the venom-containing nematocyst cells, confirming their toxin status. These findings emphasize the critical importance of managing CoTS outbreaks on reefs, especially reefs with high *Acropora* sp. abundance and developing active management strategies to reduce the impacts of CoTS.

## Data accessibility

Raw RNAseq sequences can be found at NCBI under the BioProject accession: PRJNA1230057. Code, scripts and data analysis can be found in the GitHub repository https://github.com/lmgorman/CoTS-RNAseq.

## Acknowledgements and funding sources

This research was supported by the Pacific Funds grant (HC/1674/CAB CoTS-PACIFIQUE) awarded to **SCM**, **HMP**, **LMG** and the ATER/CORAIL Laboratoire d’Excellence postdoctoral fellowship grants for “ AcantCorVenin Grant” awarded to **SCM** for **LMG**. **HMP** was also supported by NSF OCE 2348674.

As guests, we honor and acknowledge the Mā’ohi community, we give thanks for the land and water resources of Polynesia and the traditional custodians of the land on which this experimental work was conducted on the island of Mo’orea. Māuruuru roa. With respect to the spelling of Tahitian words, we endeavored to follow the Te Fare Vānaʻa, transcription system that is adhered to by a large segment of the Tahitian community, but also recognize other community members follow the Raapoto transcription system where the island name of Mo’orea is, for example, spelled without the ‘eta (i.e., Mo’orea).

We would also like to thank Zoe Storm and the technicians at CRIOBE - Alexandre Merciere, Yann Lacube, Benoit Espiau and Clement Castrec for fieldwork assistance and Benoit Espiau for permits and storage facilities.

All coral collections were conducted under permits issued by the French Polynesian authorities (Arrêté n° 3091/MPR/DRM and CITES n° FR1998700269-E), within the framework of a LabEx CORAIL grant “AcantCorVenin” to SCM & LG and Pacific Funds grants “COTS Pacifique” to SCM, LG, MB & HP. Ethical permits were granted by CNRS Animal Experimentation, R-13-CNRS-F1-16 to Yann Lacube and ANZCCART ComPass Animal Welfare Training certificate to SCM.

