## Supplemental File 1 for "Contrasting defensive strategies underlie differential susceptibility of corals to crown-of-thorns sea star (CoTS; *Acanthaster* cf. *solaris*) predation"

*Supplemental Information*

1. *Metalloproteinases*

Originally we observed a large diversity in the domain architecture of metalloproteinase genes containing the astacin domain, with many harbouring other protein domains and tandem repeats (**Table S6**). Metalloproteinases carry out other functions in non-venomous tissue e.g., cell to cell interactions, tissue regeneration and repair, shedding, development and immunity and these functions are associated with certain protein domains e.g., Kringle, MAM, fibronectin II, CUB, TSP1, PLAT, EGF, UMOD/GP2/OIT3-like, D8C domain (**Table S3**; Patthy et al., 1984; Engel, 1989; Bork & Beckmann, 1993; Bateman & Sandford,1999; Chen et al., 2000; Cismasiu et al., 2004; Stsiapanava et al., 2022). Thus, as the samples were taken during a CoTS attack, where corals would have experienced extensive tissue and cellular degradation, some metalloproteinases likely harboured non-venom function, and instead carried out other metalloproteinase functions e.g., extracellular matrix remodelling for tissue repair. However, deciphering venomous functions from other cellular metalloproteinase functions without functional studies should be interpreted with caution due to many non-venomous and venomous metalloproteinases containing similar protein domains e.g., metalloproteinases with TSP1, lectin C, F5_F8_type_C, EGF, Fxa_inhibition and PLAT domains were found in the nematocyst proteome of the sea anemones (Delgado et al., 2022; Barroso et al., 2025). Indeed, in jellyfish metalloproteinases have been shown to also have crucial roles in both development and venom toxicity (Gomis-Rüth & Stöcker, 2023; Liu et al., 2025). Therefore, metalloproteinases with structural similarity to other cnidarian toxic metalloproteinases were highlighted for further discussion (**Table S6**). This left 2 and 7 putative metalloproteinase toxin candidates for *A. hyacinthus* and *Porites* sp. respectively that showed structural similarity to Nematocyst expressed protein (NEP) 6 from *Nematostella vectensis* (**Table S6**). None of these putative metalloproteinases that showed structural homology with NEP6 were differentially expressed in attacked *A. hyacinthus* colonies (**Table 1, S6**). These structural homologs from *A. hyacinthus* and *Porites* sp. harboured either the astacin domain, astacin and Shk domain, astacin and TSP1_ADAMTS domain, the astacin and TSP1_ADAMTS and lectin C domains or the astacin and D8C_UMOD domains (**Table S6**). Snake venom metalloproteases (SVMPs) have been shown to bond to the C-type lectin-like proteins post-translationally during venom production and their bioactivity involves activating factor X (Fox & Serrano, 2005). Factor X forms part of the prothrombinase complex that converts prothrombin to thrombin (the key effector enzyme of the coagulation system) which is essential for the blood clotting pathway (Dahlbäck, 2000). Therefore, SVMPs that harbour C-type lectin domains are some of the most potent procoagulants (Morita, 2005). Whether this bioactivity is also present in cnidarian metalloproteinases harbouring the C-type lectin domain is unknown.

Additionally, we also found another metalloproteinase architecture containing a protein domain possibly involved in blood coagulation that is antagonist to Factor X - the Factor Xa_inhibition domain (**Table S6**). Thus, proteins harbouring the Fxa_inhibition domain can target the coagulation factor FXa in the blood and cause anticoagulation. Indeed, in snake venoms, proteases can both activate and inhibit Factor X (Mladic *et al.*, 2016). The Fxa_inhibition domain has also been found in proteins in other cnidarians such as *Nematostella,* as well as the nematocysts of sea anemones (Babonis et al., 2019; Delgado et al., 2022). Whether the metalloproteinases show similar functions in scleractinian venoms warrants future investigation.

Nematocyst expressed metalloproteinases, including NEP6, from *N. vectensis* have also shown this structural homology to Tolloid/Bone Morphogenetic Protein 1 but lack the canonical CUB domains (Moran et al., 2013). Indeed, some of the other putative metalloproteinases found in *A. hyacinthus* and *Porites* sp. showed structural homology to Blastula protease 10 from *Paracentrotus lividus* whose catalytic domain groups with Tolloid and Bone Morphogenetic Protein 1, forming a sub-family (**Table S6**; Lhomond et al., 1996). Whilst several of these genes did harbour CUB domains like true Tolloid and Bone Morphogenetic Protein 1, many showed an absence of CUB domains (**Table S6**), agreeing with NEP6-like proteins found in cnidarians. Several of the other putative metalloproteases showed structural homology to Zinc metalloproteinase nas-4 from *Caenorhabditis elegans* (**Table S6**).

Interestingly, we found several metalloproteinases in both *A. hyacinthus* and *Porites* sp. that harboured the astacin domain and either single or repeated domains of the Shk neurotoxin, ranging from one Shk domain to nine in *Porites* sp. gene *Peve_00038735* (**Table S6**). Despite not having structural similarity to NEP6 these metalloproteases may still function in envenomation, with venom-related proteins from the cnidarian *Oulactis* sp. also containing up to nine Shk repeats (Mitchell et al., 2020). Shk is a potent neurotoxin targeting potassium KV1.3 channels that was originally isolated from the sea anemone *Stichodactyla helianthus* (Castañeda et al., 1995; Pennington et al., 1995). In *Nematostella*, Shk-like neuropeptides promoted contraction in the tentacles and were toxic to vertebrates (Sachkova *et al.*, 2020). Metalloproteinases forming a heterodimeric sequence with Shk toxins may facilitate the spread of this potassium channel blocking toxin (Gerdol et al., 2019; Martín-Galiano & Sotillo, 2022). Indeed, metalloproteinases with Shk domains have been found in the venoms of other cnidarians (Ma et al., 2024; Barroso et al., 2025), with a jellyfish metalloproteinase harbouring two Shk domains (NnM469) from *Nemopilema nomurai* being shown to disrupt the cell matrix of human immortalized epidermal cells *in vitro* (Ma et al., 2024). Interestingly, the metalloproteinase with nine Shk repeats in *Porites* sp. (*Peve_00038735*) also harboured a domain common in blood coagulation factors, the discoidin (F5_F8_type_C) domain. Proteins harbouring the discoidin are mainly localised to the nematocysts in *Actinia fragacea* (Barroso et al., 2025). This same protein showed structural similarity to Tolloid without harbouring the CUB domain (**Table S6**), similar to nematocyst expressed metalloproteinases (Moran et al., 2013).

As metalloproteinases have several functions and this study gathered metalloproteinases from corals that would have had their structure remodelled during tissue destruction by CoTS, future studies will need to conduct functional characterisation to clarify which ones are involved in repair of tissue versus which ones are specifically involved in envenomation. Nonetheless, the metalloproteinases containing domains of known toxins, e.g., the Shk domain, as well as those showing structural homology to NEP6 are strong candidates.

2. *Kunitz-type neurotoxins*

Similarly to metalloproteinases, genes harboring the kunitz-type BPTI domain were only highlighted for further discussion if they showed structural similarity to other kunitz-type neurotoxins (**Table S7**). This left seven putative kunitz-type toxins with strong candidacy in *Porites* sp. and three in *A. hyacinthus* (**Table S7**) at 126-675 aa full-length, that all showed sequence homology to the kunitz toxin PcKuz1 and PcKuz3 in the zoanthid *Palythoa caribaeorum* (Liao et al., 2018; **Tables 1, S2 & S3**). Moreover, when we searched these amino acid sequences against the AlphaFold database, high structural hits to venomous snake toxins were returned e.g., Mambaquaretin-2. Interestingly, one of these proteins in *A. hyacinthus* (*Ahyacinthus14557)* was also one of the differentially expressed genes in *A. hyacinthus* colonies being attacked by CoTS (**Table S2)**. Both *Porites* sp. and *A. hyacinthus* harboured one putative kunitz-type neurotoxin that showed the highest structural similarity to the thrombin inhibitor Hemalin from *Haemaphysalis longicornis* (pLDDT = 91.38; **Table S7**)*.*

3. *CFXs*

We explored these putative CFX-like toxins (*Peve_00009963* and *Peve_00015288*) further by investigating their predicted 3D protein structure. Alphafold showed that *Peve_00015288* showed the highest structural similarity to *Chironex fleckeri* CfTX-2 (**Table S3**) which is unsurprising since the *Porites* CFX homologs in our previous study clustered alongside the JFT-1b type I CFXs - including CfTX-2 (Gorman et al., 2026). However, sequence homology, Foldseek and DALI analysis showed that *Peve_00015288* had the highest structural similarity to *Chironex fleckeri* CfTX-A - a type II JFT-1b (**Table S3, S8; Fig S1**). Contrastingly, *Peve_00009963* showed the highest structural homology to an uncharacterised protein in *Exaiptasia diaphana* (pLDDT = 80.19) and all the structurally similar hits were from unreviewed sequences (**Table S3**). *Peve_00009963* harbours the Pesticidal crystal protein, an N-terminal domain superfamily (IPR036716) that characterizes proteins whose N-terminal forms a pore (Grochulski et al., 1995), similar to PFTs and CFXs. When we searched the constructed *Peve_00009963* from CoLab against FoldSeek, the protein with the highest structural similarity from SwissProt was Toxin CaTX-A from *Alatina alata.* This match also agreed with data from ChimeraX and DALI, showing CaTX-A was the closest structural match closely followed by CfTX-A (**Table S8; Fig S2**). The structural matches for both *Peve_00009963* and *Peve_00015288* implies that these CFX-like toxins from *Porites* sp. may be more similar to type II JFT-1b toxins, contrasting with homologs found in *P. rus*, *P. compressa* and *P. australiensis* that are grouped with type I JFT-1b toxins (Gorman et al., 2026).

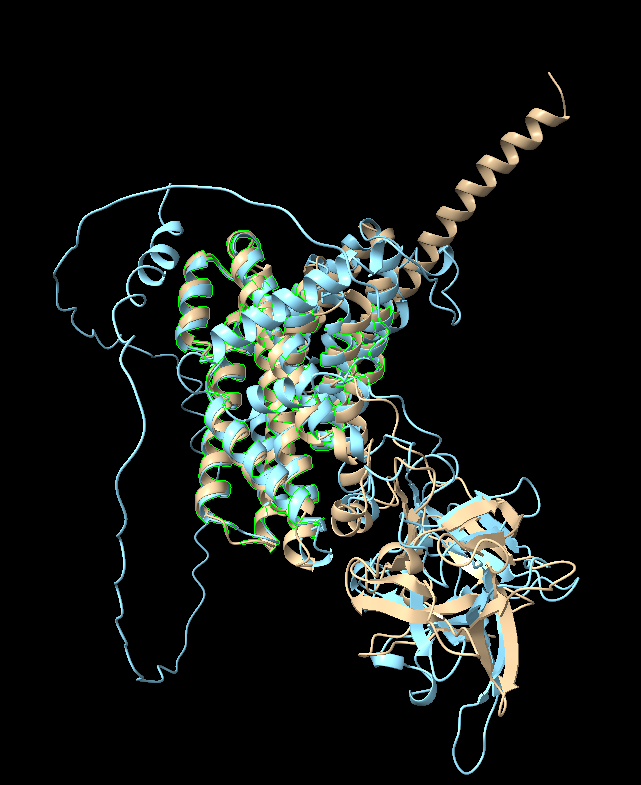

**Figure S1.** Conserved structures between *Peve_00015288* and *Chironex fleckeri* CfTX-A **B.** Putative *Porites* sp. CFX-like toxin (*Peve_00015288*) predicted structure (blue) created in ColabFold (Mirdita et al., 2022) overlaid with structure of *Chironex fleckeri* CfTX-A (beige) and their conserved structural regions (green outline). Created in ChimeraX software (Meng et al., 2023).

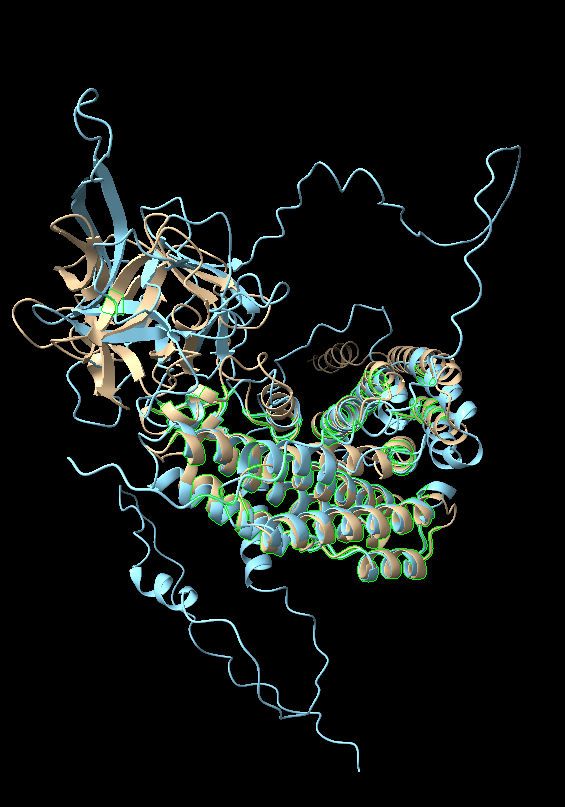

**Figure S2.** Conserved structures between *Peve_00009963* and *Alatina alata* CaTX-A **B.** Putative *Porites* sp. CFX-like toxin (*Peve_00009963*) predicted structure (blue) created in ColabFold (Mirdita et al., 2022) overlaid with structure of CaTX-A from *Alatina alata* (beige) and their conserved structural regions (green outline). Created in ChimeraX software (Meng et al., 2023).

We also investigated whether *Peve_00009963* and *Peve_00015288* harboured any regions that may represent an amphipathic helix which contributes to pore formation in PFTs (Hong et al., 2002; Malovrh et al., 2003; García-Ortega et al., 2011). The region in *Peve_00015288* that showed the strongest candidacy to an amphipathic helix occurred at residues 120-137 that had a hydrophobicity of 0.126, a hydrophobic moment of 0.466 µH and a net charge (z) of +5 (**Fig. S3**). When this region from *Peve_00015288* was aligned with *Peve_00009963* they showed conservation in their amino acid sequence and the corresponding region in *Peve_00009963* also showed a strong candidacy of forming an amphipathic helix with a hydrophobicity of 0.269, a hydrophobic moment of 0.480 µH and a net charge (z) of +5 (**Fig. S3**). A net charge (z) of 5 indicates that these parts of the proteins likely strongly interact with negatively charged surfaces e.g., cell membranes. ChimeraX also showed these regions to harbour an exposed surface (**Fig. S3**). In CFXs the transmembrane spanning region is hypothesised to also play a major role in the mode of action (Andreosso et al., 2018). Both *Peve_00009963* and *Peve_00015288* showed regions of structural conservation to CfTX-1 in the transmembrane spanning region but did not align with CfTX-1 in the amphiphilic helix region (**Fig. S4**; Andreosso et al., 2018).

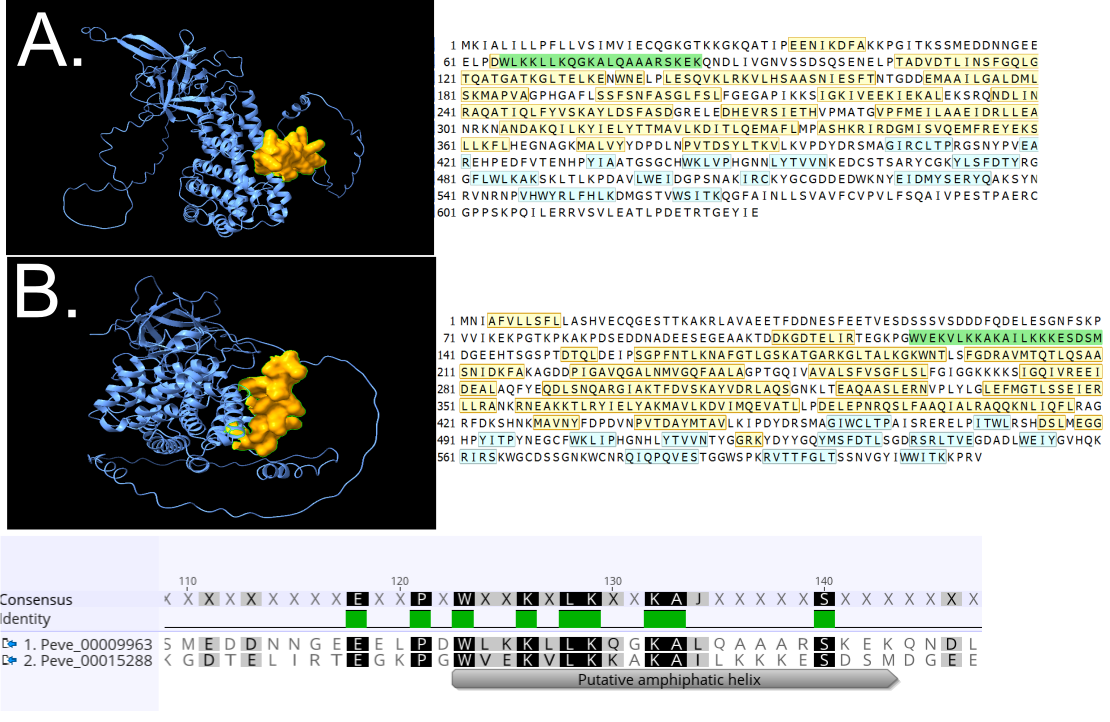

**Figure S3.** Putative CFX-like toxin sequence of *Peve_00009963* (**A**) and *Peve_00015288* (**B**) showing structural helices (yellow) and B strands (blue) with the region highlighted as a strong candidate for the amphipathic helix in green, created in ChimeraX software (Meng et al., 2023). Pairwise Sequence Alignment of putative amphiphilic helix regions of *Peve_00009963* and *Peve_00015288* shown below in Geneious Prime using the Geneious Alignment tool with Blosum 62 matrix and default parameters.

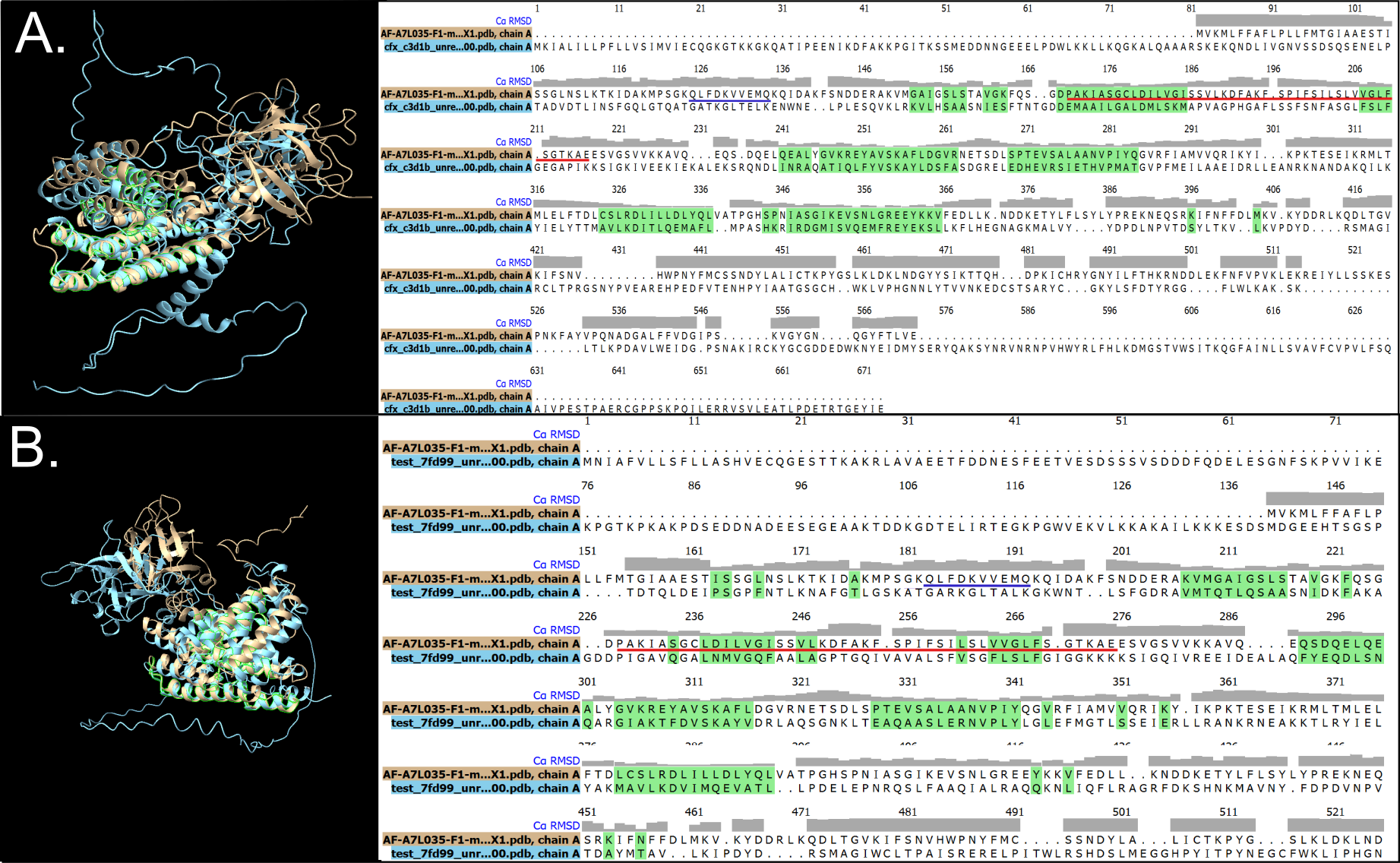

**Figure S4**. Structural alignment of *Peve_00009963* (**A**, blue) and *Peve_00015288* (**B**, blue) with *Chironex fleckeri* CfTX-1 (beige), created in ChimeraX software (Meng et al., 2023). CfTX-1 amphiphilic helix region and the transmembrane spanning region are underlined in blue and red, respectively (Andreosso et al., 2018). Areas with structural conservation are highlighted in green.

4. *Other putative venom components*

Additionally, we found genes with annotations that may correlate with a function in envenomation that were not found using searches with the UniProt ToxProt e.g., venom allergens and SE-cephalotoxin-like proteins (**Table S9**).

*References*

Andreosso, A., Bansal, P.S., Smout, M.J., Wilson, D., Seymour, J.E. and Daly, N.L., 2018. Structural characterisation of predicted helical regions in the Chironex fleckeri CfTX-1 toxin. *Marine Drugs*, *16*(6), p.201

Babonis, L.S., Ryan, J.F., Enjolras, C. and Martindale, M.Q., 2019. Genomic analysis of the tryptome reveals molecular mechanisms of gland cell evolution. *Evodevo*, *10*(1), p.23.

Barroso, R.A., Rodrigues, T., Campos, A., Almeida, D., Guardiola, F.A., Turkina, M.V. and Antunes, A., 2025. Proteomic diversity of the sea anemone Actinia fragacea: comparative analysis of nematocyst venom, mucus, and tissue-specific profiles. *Marine Drugs*, *23*(2), p.79.

Bateman, A. and Sandford, R., 1999. The PLAT domain: a new piece in the PKD1 puzzle. *Current Biology*, *9*(16), pp.R588-R590.

Bork, P. and Beckmann, G., 1993. The CUB domain: a widespread module in developmentally regulated proteins. *Journal of molecular biology*, *231*(2), pp.539-545.

Castañeda, O., Sotolongo, V., Amor, A.M., Stöcklin, R., Anderson, A.J., Harvey, A.L., Engström, Å., Wernstedt, C. and Karlsson, E., 1995. Characterization of a potassium channel toxin from the Caribbean Sea anemone Stichodactyla helianthus. *Toxicon*, *33*(5), pp.603-613.

Chen, H., Herndon, M.E. and Lawler, J., 2000. The cell biology of thrombospondin-1. *Matrix Biology*, *19*(7), pp.597-614.

Cismasiu, V.B., Denes, S.A., Reilander, H., Michel, H. and Szedlacsek, S.E., 2004. The MAM (meprin/A5-protein/PTPmu) domain is a homophilic binding site promoting the lateral dimerization of receptor-like protein-tyrosine phosphatase μ. *Journal of biological chemistry*, *279*(26), pp.26922-26931.

Dahlbäck, B., 2000. Blood coagulation. *The Lancet*, *355*(9215), pp.1627-1632.

Delgado, A., Benedict, C., Macrander, J. and Daly, M., 2022. Never, ever make an enemy… out of an anemone: transcriptomic comparison of clownfish hosting sea anemone venoms. *Marine Drugs*, *20*(12), p.730.

Engel, J., 1989. EGF-like domains in extracellular matrix proteins: localized signals for growth and differentiation?. *FEBS letters*, *251*(1-2), pp.1-7.

Fox, J.W. and Serrano, S.M., 2005. Structural considerations of the snake venom metalloproteinases, key members of the M12 reprolysin family of metalloproteinases. *Toxicon*, *45*(8), pp.969-985.

García-Ortega, L., Alegre-Cebollada, J., García-Linares, S., Bruix, M., Martínez-del-Pozo, Á. and Gavilanes, J.G., 2011. The behavior of sea anemone actinoporins at the water–membrane interface. *Biochimica et Biophysica Acta (BBA)-Biomembranes*, *1808*(9), pp.2275-2288.

Gerdol, M., Cervelli, M., Mariottini, P., Oliverio, M., Dutertre, S. and Modica, M.V., 2019. A recurrent motif: diversity and evolution of ShKT domain containing proteins in the vampire snail Cumia reticulata. *Toxins*, *11*(2), p.106.

Gomis-Rüth, F.X. and Stöcker, W., 2023. Structural and evolutionary insights into astacin metallopeptidases. Frontiers in Molecular Biosciences, 9, p.1080836.

Gorman, L.M., Huffmyer, A.S., Byrne, M., Mills, S.C. and Putnam, H.M., 2026. Coral Venom and Toxins as Protection Against Crown‐of‐Thorns Sea Star Attack. *Molecular Ecology*, p.e70202.

Grochulski, P., Masson, L., Borisova, S., Pusztai-Carey, M., Schwartz, J.L., Brousseau, R. and Cygler, M., 1995. Bacillus thuringiensisCrylA (a) insecticidal toxin: crystal structure and channel formation. *Journal of molecular biology*, *254*(3), pp.447-464.

Hong, Q., Gutiérrez-Aguirre, I., Barlic, A., Malovrh, P., Kristan, K., Podlesek, Z., Macek, P., Turk, D., González-Manas, J.M., Lakey, J.H. and Anderluh, G., 2002. Two-step membrane binding by Equinatoxin II, a pore-forming toxin from the sea anemone, involves an exposed aromatic cluster and a flexible helix. *Journal of biological chemistry*, *277*(44), pp.41916-41924.

Lhomond, G., Ghiglione, C., Lepage, T. and Gache, C., 1996. Structure of the gene encoding the sea urchin blastula protease 10 (BP10), a member of the astacin family of Zn2+–metalloproteases. *European journal of biochemistry*, *238*(3), pp.744-751.

Liao, Q., Li, S., Siu, S.W.I., Yang, B., Huang, C., Chan, J.Y.W., Morlighem, J.E.R., Wong, C.T.T., Radis-Baptista, G. and Lee, S.M.Y., 2018. Novel Kunitz-like peptides discovered in the zoanthid Palythoa caribaeorum through transcriptome sequencing. *Journal of proteome research*, *17*(2), pp.891-902.

Liu, X., Peng, X., Wang, J., Ju, S., Sun, Q., Ji, W., Hua, X., Zhang, H., Höfer, J., Pozzolini, M. and Xu, S., 2025. Targeting metzincins to mitigate jellyfish blooms: a novel approach for conservation. *Frontiers in Marine Science*, *12*, p.1563258.

Ma, Y., Yu, H., Teng, L., Geng, H., Li, R., Xing, R., Liu, S. and Li, P., 2024. NnM469, a novel recombinant jellyfish venom metalloproteinase from Nemopilema nomurai, disrupted the cell matrix. *International Journal of Biological Macromolecules*, *281*, p.136531.

Malovrh, P., Viero, G., Dalla Serra, M., Podlesek, Z., Lakey, J.H., Macek, P., Menestrina, G. and Anderluh, G., 2003. A novel mechanism of pore formation: membrane penetration by the N-terminal amphipathic region of equinatoxin. *Journal of Biological Chemistry*, *278*(25), pp.22678-22685.

Martín-Galiano, A.J. and Sotillo, J., 2022. Insights into the functional expansion of the astacin peptidase family in parasitic helminths. *International journal for parasitology*, *52*(4), pp.243-251.

Meng, E.C., Goddard, T.D., Pettersen, E.F., Couch, G.S., Pearson, Z.J., Morris, J.H. and Ferrin, T.E., 2023. UCSF ChimeraX: Tools for structure building and analysis. *Protein Science*, *32*(11), p.e4792.

Mirdita, M., Schütze, K., Moriwaki, Y., Heo, L., Ovchinnikov, S. and Steinegger, M., 2022. ColabFold: making protein folding accessible to all. *Nature methods*, *19*(6), pp.679-682.

Mitchell, M.L., Tonkin-Hill, G.Q., Morales, R.A., Purcell, A.W., Papenfuss, A.T. and Norton, R.S., 2020. Tentacle transcriptomes of the speckled anemone (Actiniaria: Actiniidae: Oulactis sp.): venom-related components and their domain structure. *Marine Biotechnology*, *22*(2), pp.207-219.

Mladic, M., Zietek, B.M., Iyer, J.K., Hermarij, P., Niessen, W.M., Somsen, G.W., Kini, R.M. and Kool, J., 2016. At-line nanofractionation with parallel mass spectrometry and bioactivity assessment for the rapid screening of thrombin and factor Xa inhibitors in snake venoms. *Toxicon*, *110*, pp.79-89.

Moran, Y., Praher, D., Schlesinger, A., Ayalon, A., Tal, Y. and Technau, U., 2013. Analysis of soluble protein contents from the nematocysts of a model sea anemone sheds light on venom evolution. *Marine biotechnology*, *15*(3), pp.329-339.

Morita, T., 2005. Structures and functions of snake venom CLPs (C-type lectin-like proteins) with anticoagulant-, procoagulant-, and platelet-modulating activities. *Toxicon*, *45*(8), pp.1099-1114.

Patthy, L., Trexler, M., Váli, Z., Banyai, L. and Varadi, A., 1984. Kringles: modules specialized for protein binding: homology of the gelatin-binding region of fibronectin with the kringle structures of proteases. *FEBS letters*, *171*(1), pp.131-136.

Pennington, M.W., Byrnes, M.E., Zaydenberg, I., Khaytin, I., De Chastonay, J., Krafte, D.S., Hill, R., Mahnir, V.M., Volberg, W.A., Gorczyca, W. and Kem, W.R., 1995. Chemical synthesis and characterization of ShK toxin: a potent potassium channel inhibitor from a sea anemone. *International journal of peptide and protein research*, *46*(5), pp.354-358.

Sachkova, M.Y., Landau, M., Surm, J.M., Macrander, J., Singer, S.A., Reitzel, A.M. and Moran, Y., 2020. Toxin-like neuropeptides in the sea anemone Nematostella unravel recruitment from the nervous system to venom. *Proceedings of The National Academy of Sciences*, *117*(44), pp.27481-27492.

Stsiapanava, A., Xu, C., Nishio, S., Han, L., Yamakawa, N., Carroni, M., Tunyasuvunakool, K., Jumper, J., de Sanctis, D., Wu, B. and Jovine, L., 2022. Structure of the decoy module of human glycoprotein 2 and uromodulin and its interaction with bacterial adhesin FimH. *Nature Structural & Molecular Biology*, *29*(3), pp.190-193.
